## Supplementary Figures for "Cell type-dependent differential activation of ERK by oncogenic KRAS in colon cancer and intestinal epithelium" (from another version of this preprint: version 2, 2019-02-13)

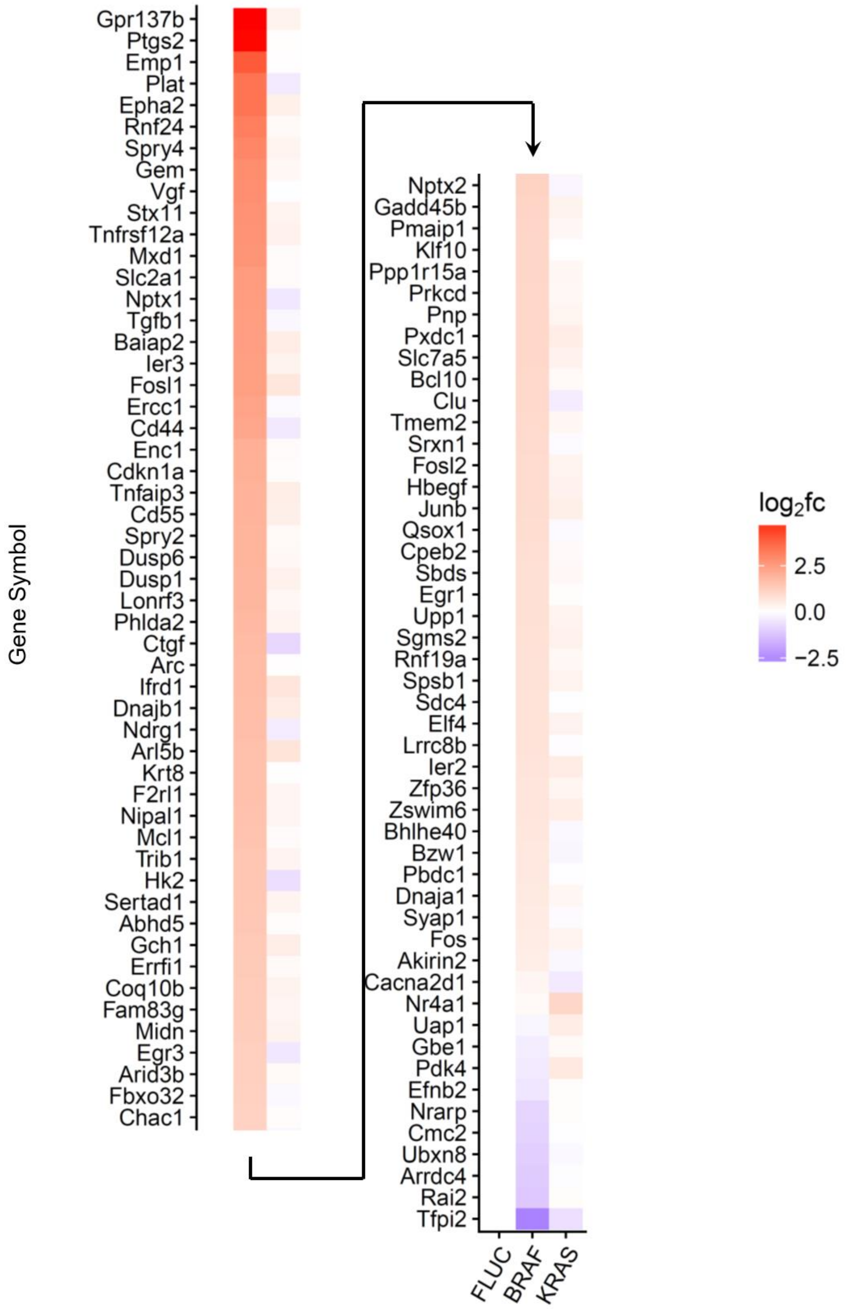


**Figure S1: Bulk tissue transcriptome analysis reveals near-universal upregulation of MAPK targets by BRAF$^{V600E}$.** MAP kinase target genes, as defined in [34] were assessed in transcriptomes 24 h after transgene induction. Upregulation was found after BRAF$^{V600E}$, but not after KRAS$^{G12V}$, induction. Replicate transcriptomes were generated from transgene-positive cells of induced organoids, as described before [58]. RNA-seq data are available in under accession number GSE93947.

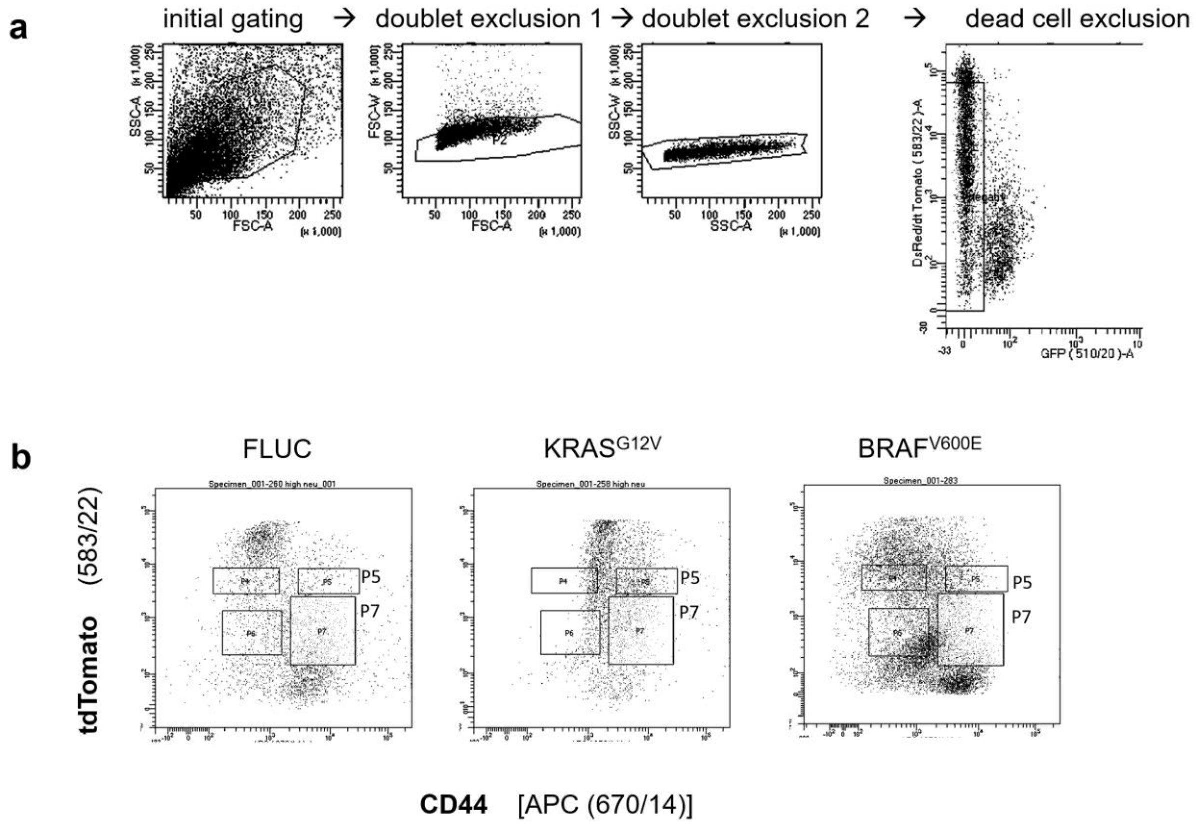


**Figure S2: FACS gating strategy for FLUC-, $KRAS^{G12V}$-, and $BRAF^{V600E}$ cells for single cell sequencing.** a) Gating strategy to obtain alive single cells. b) Representative example of tdTomato (transgene; 583/22 bandpass (BP) filter) and CD44 (670/14 BP filter) fluorescence in transgene-induced organoid cell suspensions, as indicated. Note that induction of $KRAS^{G12V}$, but not of $BRAF^{V600E}$, increases levels of crypt cell marker CD44. This effect is likewise seen in the CyTOF data sets (compare Fig. 6E). Gate P5 was used to sort $KRAS^{G12V}$-induced cells. Gate P7 was used to sort for $BRAF^{V600E}$-induced cells. The alternative gate (as compared to $KRAS^{G12V}$ cells) was used, as high $BRAF^{V600E}$ expression results in intestinal cell death (see Fig. 2). Both gates, P5 and P7, were employed for FLUC-induced cells.

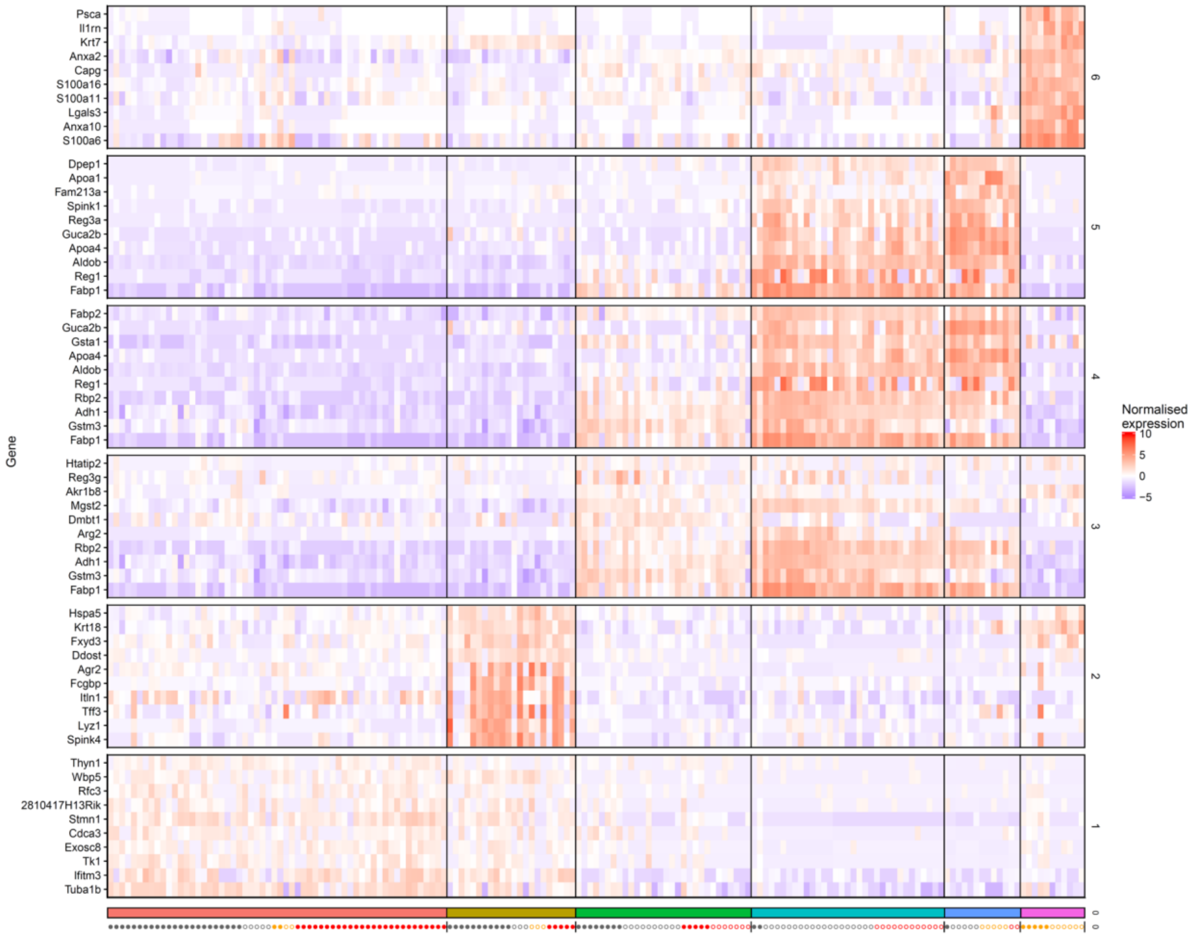


**Figure S3: Signature genes for six transcriptome clusters derived from FLUC, KRAS$^{G12V}$ and BRAF$^{V600E}$-induced organoid cells.** Top 10 overrepresented genes per cluster are given (bottom to top).

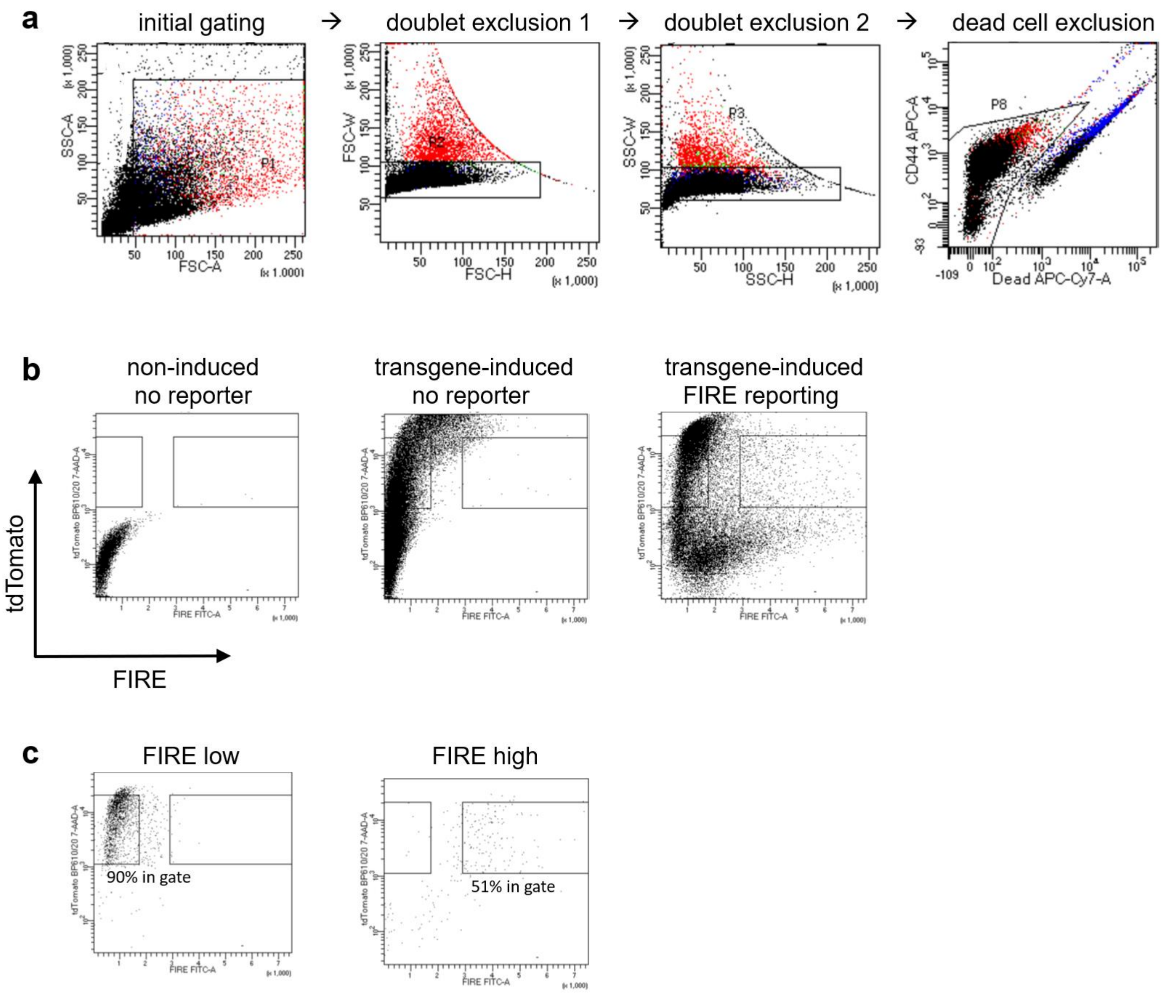


**Figure S4: FACS gating strategy for transgene induced FIRE-positive and -negative cells for single cell sequencing.** a) Gating strategy to obtain alive single cells b) Representative example of tdTomato (transgene; 610/20 BP) and FIRE (ERK reporter; 530/30 BP) fluorescence in control and transgenic organoid cell suspensions, as indicated. FIRE fluorescence is low and detected in linear detector mode c) Post sort analysis for FIRE-positive and negative cells. In short, induced organoid cells were sorted into BSA-coated tubes, using FIRE-negative and FIRE-positive gates, as indicated. Numbers indicate percent of target population from all events.

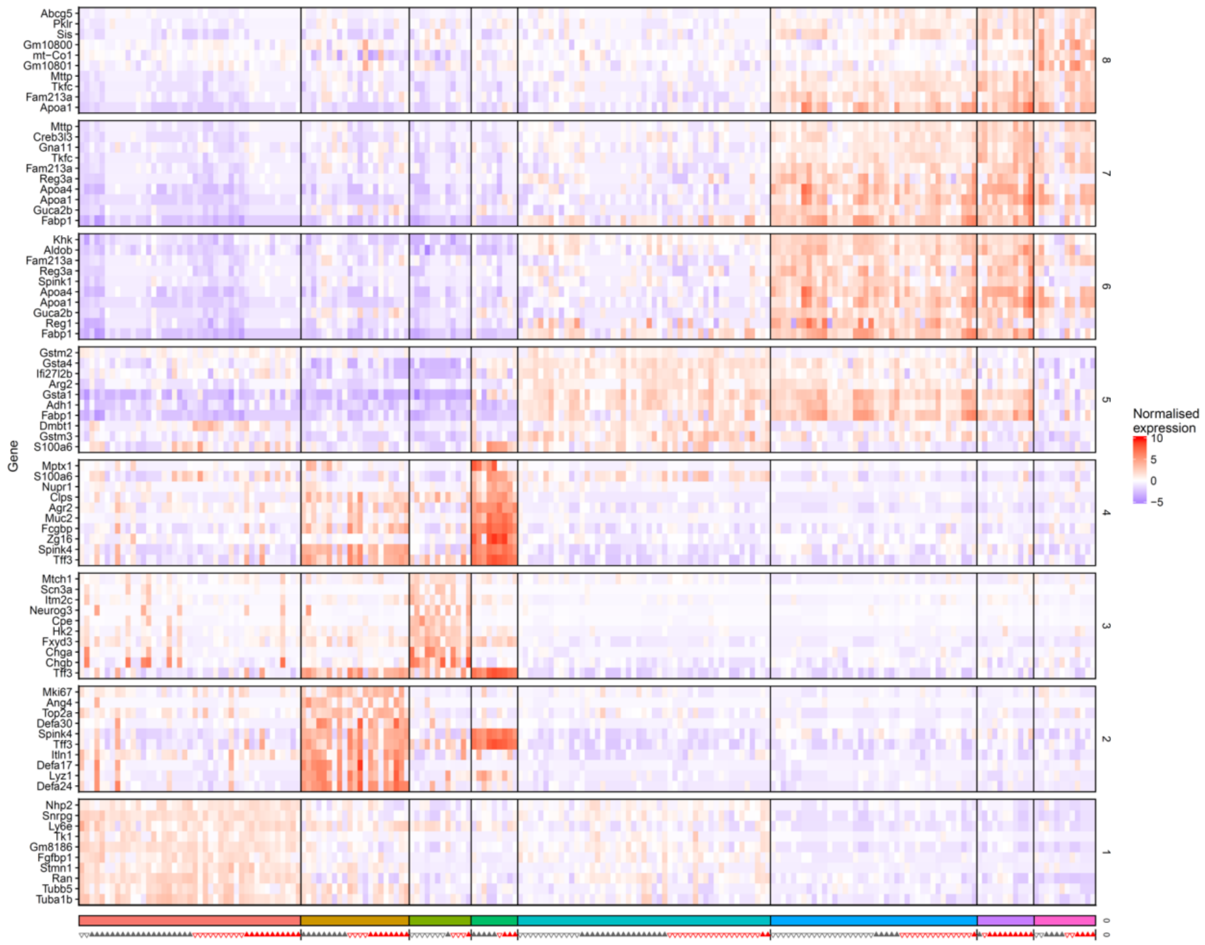

**Figure S5: Signature genes for eight transcriptome clusters derived from organoid cells sorted by FIRE activity.** Top 10 overrepresented genes per cluster are given (bottom to top).

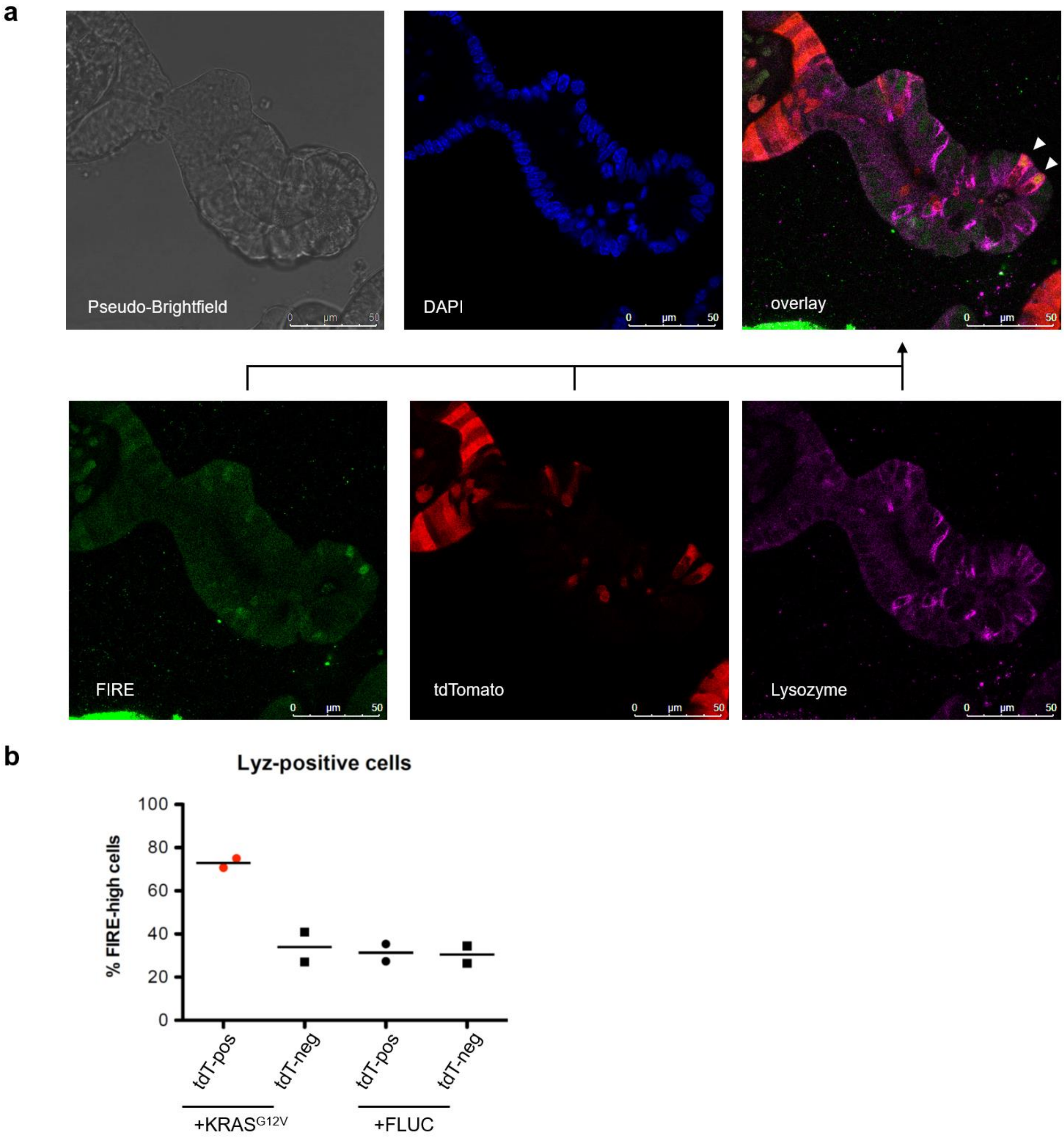


**Figure S6: Immunofluorescence confirms Paneth cells as a main FIRE-positive cell type in the crypt**. a) Pseudo brightfield image along with fluorescent channels representing nuclear stain (DAPI; blue), FIRE fluorescence (Venus; green), transgene fluorescence (tdTomato; red), and Paneth cell immunostaining (Lysozyme; purple) are given. Overlay combines the green, red and purple channels. b) Quantification of FIRE-high Lysozyme (Lyz)-positive Paneth cells after induction of KRAS$^{G12V}$- or FLUC-encoding transgenes. Data represents 2 independent experiments with >100 cells Lyz(+) cells counted per condition.

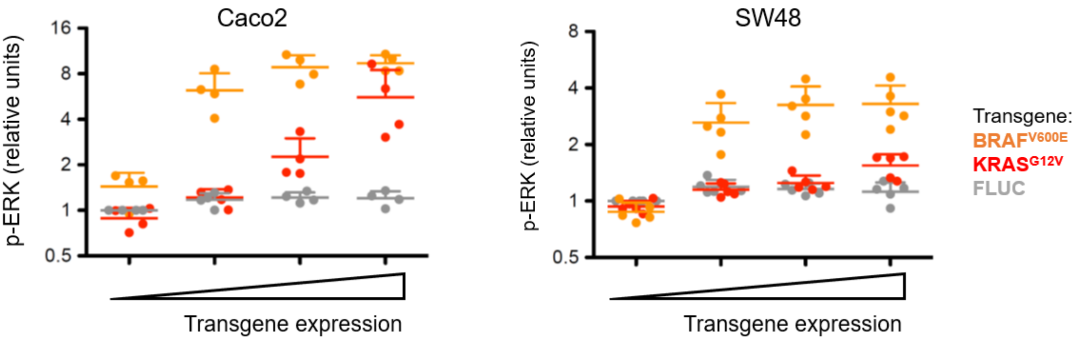


**Figure S7: Signal transduction from KRAS$^{G12V}$ to ERK differs between CRC cell lines** Quantification of phospho-ERK in KRAS$^{G12V}$-, BRAF$^{V600E}$- or FLUC-transfected Caco2 and SW48 CRC cells, by flow cytometry. Mean and standard deviation of 4 independent replicate experiments is given. Grey: FLUC-; orange: BRAF$^{V600E}$; red: KRAS$^{G12V}$-transfected cells. SW48 and Caco2 CRC cell lines have no mutations in KRAS, NRAS or BRAF. For the experiments, cells were transfected with inducible expression vectors for tdTomato-KRAS$^{G12V}$, -BRAF$^{V600E}$, or -FLUC control, as previously used in organoid cultures. 24 h after induction, cells were fixed and oncogene levels, as extrapolated from tdTomato, and phosphorylation of ERK were measured via flow cytometry. We found that even low levels of BRAF$^{V600E}$ (orange dots) strongly increased ERK phosphorylation in both cell lines. In contrast, the ability of KRAS$^{G12V}$ (red dots) to induce ERK phosphorylation was markedly different. In Caco2 cells, ERK phosphorylation increased stepwisely with KRAS$^{G12V}$-associated fluorescence, whereas in SW48 even high levels of transgenic tdTomato-KRAS$^{G12V}$ were unable to induce strong ERK phosphorylation.

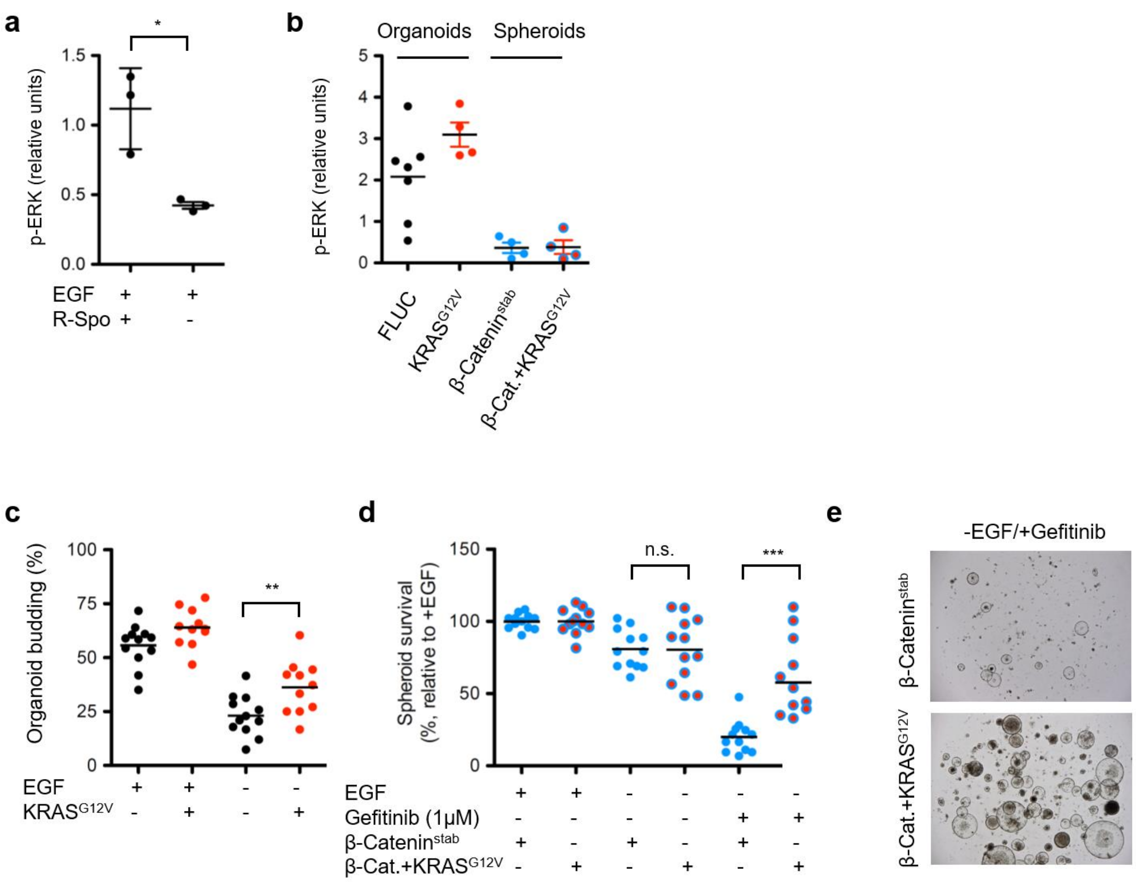


**Figure S8: Interaction between β-Catenin and RAS-ERK signalling in intestinal organoids**. a) p-ERK quantification by capillary protein analysis, 24 h after R-Spondin removal. Results indicate decrease of ERK phosphorylation after abrogation of Wnt/β-Catenin activity in organoids. b) phospho-ERK quantification by capillary protein analysis in spheroid cells that have undergone a switch to an adenoma-like phenotype after long-term activation of stabilized β-Catenin and removal of the Wnt co-ligand R-Spondin. The results indicate decreasing ERK phosphorylation in cultures with constitutively high β-Catenin activity. We propose that the changes of p-ERK levels observed in a) and b) are due to shifts in cell type. c)-e) Functional interaction between $KRAS^{G12V}$ and β-Catenin. c) Quantification of organoid budding 4 days after induction of oncogenic $KRAS^{G12V}$ in the presence or absence of EGF in normal tissue organoids. Cysts forming after organoid disaggregation were counted 24 h after passaging. Fractions of budding organoids (that is, of cysts that developed a crypt domain with a distinct adjacent villus domain, as indicated by an oblong form) were calculated at day 4. It is of note that although a significant increase in budding cysts was observed, organoids could never be propagated after passaging in the absence of EGF, not even after induction of $KRAS^{G12V}$. d) Quantification of spheroid survival after EGF depletion, EGFR inhibition by Gefitinib and/or $KRAS^{G12V}$ induction, 6 days after passaging. Quantification from multiple wells of 3 independent experiments e) Representative images of β-$Catenin^{stab}$- or β -$Catenin^{stab}$/$KRAS^{G12V}$ double transgenic spheroid cultures, 6 days after passaging and in EGF-free medium in the presence of 1 μM Gefitinib.
